# Determination of clinical risk factors associated with inflammation in hypertensive patients with type-2 diabetes mellitus

**DOI:** 10.1101/613711

**Authors:** Mohammed S. Ellulu

## Abstract

**Background:** Obesity and chronic diseases associated with the development of inflammation have remained unclear if the observed inflammatory state in diabetic patients is due to excess adipose tissue mass and/or directly associated with the diabetic state. Therefore, this study determined the risk factors associated with inflammation in hypertensive patients with type-2 diabetes mellitus.

**Methods:** A total of 164 hypertensive diabetic patients aged 38 to 60 years were selected from seven primary health care centers in Gaza city, Palestine. Interview and questionnaire were employed to collect data related to age, gender, smoking habits, and physical activity pattern. Besides, the selection of patients depended on objective criteria.

**Results:** The study involved 118 (72%) women and 46 (28%) men. The mean of age for all patients was 53.7±0.46 years old. 76 patients (46.3%) were categorized as current smokers, 88 patients (53.7%) categorized as non-smokers. The baseline distribution of patients according to physical activity has displayed that 130 (79.3%) were low physically active patients, 28 (27.1%) were moderate, and 6 (3.7%) were highly physically active patients. A tertile of inflammation feature with high sensitivity C-reactive protein (hs-CRP) was developed. The highest tertile of hs-CRP was significantly associated with women, higher obesity indices, metabolic dysregulation involving lipid profile markers, fasting blood glucose (FBG) and blood pressure, higher interleukin 6 (IL-6), and lower adiponectin. Via ordinal logistic regression analysis, after adjusting for age, gender, smoking habits, and physical activity; the risk factors for hs-CRP were the increased body mass index [OR: 1.17, P=0.018], IL-6 [OR: 2.22, P=0.025] and FBG [OR: 1.01, P=0.007], as well as reduced adiponectin [OR: 0.81, P=0.002].

**Conclusion:** The inflammation state was affected by obesity and had been related to altered adipokines levels of IL-6 and adiponectin, as well as affected by the disease condition of diabetes, as evidenced by higher serum level of FBG.

## INTRODUCTION

The World Health Organization (WHO) rates hypertension (HT) as one of the most important causes of premature death worldwide [1]. It is one of the most important preventable causes of death worldwide and one of the commonest conditions treated in primary care in the United Kingdom, where it affects more than a quarter of all adults and over half of those over the age of 65 [2]. In addition, HT was ranked as the third most important risk factor for attributable burden of disease in south Asia in 2010 [3]. In fact, the uncontrolled HT has been associated with developed nephropathy, retinopathy, ischemic heart disease, peripheral vascular disease [4], and inflammation [5,6]. According to the German Institute of Human Nutrition *(DIFE),* the uncontrolled HT has been associated with the development of type-2 diabetes mellitus (T2DM) [7].

Diabetes mellitus is a group of diseases marked by high level of blood glucose [8]. According to the *American Diabetes Association,* the T2DM is a result of insulin resistance that leads to a progressive defect of insulin secretion [9]. T2DM leads to progressive complications like neuropathy, retinopathy, kidney disease, and heart disease [10]. It also worsens arterial stiffness in hypertensive patients through endothelial dysfunction [11], and increases the incidence of metabolic dysregulation, oxidative stress, and inflammation [12,13]. From these associated complications; patients suffer greatly, and result in an increased number of years lived with disability [14].

On the other hand, obesity, the accumulation of abnormal or excessive fat [15], is the fifth leading risk for global deaths [16]. It has been classified as one of the primary causes of elevated blood pressure (BP) among adults [17], and increases the risk of developing HT irrespective of race or gender [18]. Furthermore, obesity, and especially visceral adipose tissue accumulation, increases the risk of impaired insulin tolerance [19] and the development of T2DM [20]. The greater risk of T2DM among the obese can partly be explained by the changes in adipose tissue function [21]. Besides, epidemiological studies have demonstrated inflammation evidenced by the elevated plasma levels of inflammatory markers, including C-reactive protein (CRP), interleukin 6 (IL-6), and tumor necrosis factor alpha (TNF-α), in patients with metabolic syndrome and T2DM [22,23]. Moreover, the studies have indicated lower plasma level of adiponectin, which is totally antiinflammatory markers and insulin-sensitizing factors, in obese individuals [24], cardiovascular patients, metabolic syndrome, and T2DM [25].

As reported, the conditions of obesity, HT, and T2DM are well associated with inflammation [26,27]. Thence, inflammation due to increased plasma level of CRP can lead, over the time, to metabolic syndrome [28], atherosclerosis and coagulation [29], cardio-metabolic risks [30], and heart attack [31]. However, it remains unclear if the observed alterations in plasma cytokines and/or inflammatory markers in T2DM are due to excess adipose tissue mass and/or directly associated with the diabetic state [32]. Therefore, this study identified the clinical risk factors associated with inflammation evidenced by high sensitivity C-reactive protein (hs-CRP) in hypertensive diabetic patients.

## RESEARCH DESIGN AND METHODS

### Study subjects

In this study, a total of 164 hypertensive patients with type-2 diabetes mellitus *“hypertensive diabetic*” out of 484 subjects screened; those non-insulin dependent and aged 38-60 years old were selected from seven primary health care centers that were controlled by the Ministry of Health in Gaza city, Palestine via cluster random sampling, from November 2013 to May 2014. The study was ethically approved by the Helsinki Committee for Ethical Approval of Gaza (Number: PHRC/HC/11/13). All patients had a stable weight with no fluctuation of >2% of their body weight for at least two months prior to this study. Other than that, patients who suffered conditions that might affect the laboratory results like acute or chronic inflammatory diseases, including infections, arthritis, cancers, heart disease, liver and renal diseases, or regularly used medicines like insulin, NSAIDs such as COX-2 inhibitors, and antibiotics, were excluded. Furthermore, physiological changes like pregnancy and breast feeding states were also excluded.

Personal information collected via interview and questionnaire involved age, gender, health/disease history, and medicine usage. Besides, the lifestyle factors involved smoking habits and physical activity pattern. The smoking habits were assessed based on modified form of The Behavioral Risk Factor Surveillance System that had been approved by the Centers for Disease Control and Prevention (CDC) Survey Data [33]. The patients were categorized into two groups; the first group comprised of current smokers that involved both active and passive smokers, while the second group consisted of non-smokers that involved past smokers and never smokers. Meanwhile, physical activity pattern was evaluated based on the Global Physical Activity Questionnaire (GPAQ) Version-2 that had considered the Palestinian specialties [34]. The patients were grouped into one out of the three categories based on the intensity of physical activity that had been classified as low, moderate, and high.

### Tools

A Seca Stadiometer was used to assess body mass index (BMI) based on the World Health Organization, calculated by using height and weight based on the formula [BMI (kg/m^2^) = Weight (kg) / Height square (m^2^)] [35], and a Seca 201 non-elastic tape was used to determine WC based on the National Institute for Health and Clinical Excellence classification [36] (men: normal <102 cm, high ≥102 cm; women: normal <88 cm, high ≥88 cm).

Besides, quantitative method was employed to assess fasting blood glucose (FBG), total cholesterol (TC), triglyceride (TG), and high sensitivity C-reactive protein (hs-CRP). A CRP turbidimetric latex 1:5 kit was used to assess hs-CRP, while an enzymatic colorimetric method with glucose oxidase was used to estimate FBG, and commercial kits were used to assess TC and TG. All quantitative data were evaluated by using a Mindray BS-120 Chemistry Analyzer. Meanwhile, interleukin 6 (IL-6) and adiponectin were assessed by using enzyme-linked immunosorbent assay (ELISA) kits (Sigma-Aldrich Co) via One-Run reading. Besides, blood samples were collected by trained nurses or physicians at the health centers. The 7 mL blood sample was drawn into a polyethylene evacuated tube, and then, divided into two separate tubes. One of the tubes that contained blood sample was used to evaluate quantitative biochemical measures (hs-CRP, FBG, TC, and TG), while the other was stored at −80°C, after separation from serum, for analysis through one-run of an ELISA reader to assess IL-6 and adiponectin.

In addition, BP was determined through validation and standard mercury sphygmomanometer, where systolic BP (SBP) was defined as the appearance of first sound (Korotkoff phase 1), and diastolic BP (DBP) was the disappearance of the sound (Korotkoff phase 5) during deflation of the cuff at a 2-3 mmHg/second reduction rate of the mercury column. Two successive BP readings were obtained at 5 minutes intervals and averaged.

### Statistical analysis

The data obtained were analyzed by using the Statistical Package for Social Sciences version 21.0 software (SPSS Inc., Chicago, IL, USA). Descriptive statistics, including Chi-square (χ^2^), was used to compare the categorical variables. The central tendency of continuous variables was presented by mean ± standard error of mean (SEM). Then, a tertile of hs-CRP was created; where One-Way ANOVA was used to detect the differences between the tertile groups, whereas the Ordinal Logistic Regression was used to estimate the Exp(B) [Odds Ratio (OR)] after adjustments were made for age, gender, smoking habits, and physical activity. Meanwhile, the Bivariate analysis via Pearson’s correlation coefficient explained the relationships of hs-CRP with BMI, IL-6, adiponectin, and FBG. The p value of ≤0.05 was considered as statistically significant and the level of confidence was 95%.

## RESULTS

A total of 164 HT patients with T2DM were recruited in this study to identify the clinical risk factors associated with inflammation evidenced by hs-CRP. The study involved 118 (72%) women and 46 (28%) men. The mean of age for all patients was 53.7±0.46 years old. Lifestyle habits included smoking and physical activity patterns. 76 patients (46.3%) who were categorized as current smokers had been active and passive smokers, while 88 patients (53.7%) categorized as non-smokers were past smokers and never smokers. Finally, the baseline distribution of patients, according to physical activity, displayed high percentage of low physical activity for 130 patients (79.3%), 28 (27.1%) were moderate, and 6 (3.7%) were highly physically active patients.

As shown in **Table 1**, the recruited patients were distributed into one of three groups based on hs-CRP tertile. The tertile of hs-CRP (mg/L) was categorized as in the following: Low, hs-CRP<3.71 mg/L (n=55); Moderate, hs-CRP=3.71-8.99 mg/L (n=54); and High, hs-CRP>8.99 mg/L (n=55). The distribution of gender to hs-CRP tertile revealed significant difference, as the percentages of women increased high-moderate-low tertile movement, while the percentages of men moved inversely; they decreased in high-moderate-low tertile movement. However, no difference was obtained for age due to the changed hs-CRP tertile, as well as the changes of smoking habits and physical activity that exhibited non-significant distribution.

**Table 1:** Distribution of participants’ characteristics according to serum hs-CRP tertile]

|  | Tertile of plasma hs-CRP (mg/L) |  |  | Estimate<br>95% CI | OR/<br>Exp(B) | P<br>value |
| --- | --- | --- | --- | --- | --- | --- |
|  | Low<br>(<3.71) | Moderate<br>(3.71–8.99) | High<br>(>8.99) |  |  |  |
| Number | 55 | 54 | 55 |  |  |  |
| hs-CRP (mg/L)** | 2.16±0.13 | 6.27±0.22 | 17.77±1.88 |  |  |  |
| Age (years) | 53.85±0.79 | 53.74±0.86 | 53.49±0.76 |  |  | 0.281 |
| Gender** |  |  |  |  |  |  |
| %Women | 26.3 | 34.7 | 39.0 |  |  | 0.067 |
| %Men | 52.2 | 28.3 | 19.6 |  |  | -- |
| Smoking habits |  |  |  |  |  |  |
| %Current smokers | 36.8 | 31.6 | 31.6 |  |  | 0.665 |
| %Non-smokers | 30.7 | 34.1 | 35.2 |  |  | -- |
| Physical activity |  |  |  |  |  |  |
| %Low | 30.0 | 33.1 | 36.9 |  |  | 0.259 |
| %Moderate | 42.9 | 32.1 | 25.0 |  |  | 0.483 |
| %High | 66.7 | 33.3 | -- |  |  | -- |
| WC (cm)** | 112.62±1.67 | 117.78±1.95 | 119.95±1.49 |  |  | 0.372 |
| **BMI (kg/m<sup>2</sup>)\*\*** | 27.69±0.95 | 34.86±1.28 | 37.86±0.81 | **0.163**<br>(0.028 to 0.298) | **1.17** | **0.018** |
| SBP (mmHg) | 138.24±3.86 | 140.68±3.57 | 145.53±3.06 |  |  | 0.652 |
| DBP (mmHg)* | 80.42±2.38 | 82.84±1.84 | 88.73±1.58 |  |  | 0.117 |
| **FBG (mg/dL)\*\*** | 146.90±12.92 | 177.13±13.99 | 207.33±9.64 | **0.010**<br>(0.003 to 0.017) | **1.01** | **0.007** |
| TC (mg/dL)** | 194.10±4.95 | 208.03±7.28 | 209.69±5.37 |  |  | 0.390 |
| TG (mg/dL)** | 166.76±18.08 | 234.10±24.00 | 235.69±13.76 |  |  | 0.640 |
| **IL-6 (pg/mL)\*\*** | 1.60±0.10 | 2.05±0.12 | 2.39±0.11 | **0.802**<br>(0.102 to 1.502) | **2.22** | **0.025** |
| **Adiponectin (mg/L)\*\*** | 13.20±0.98 | 10.93±0.62 | 8.55±0.53 | **-0.204**<br>(-0.333 to -0.076) | **0.81** | **0.002** |

Central tendency measured by Mean ± SEM.
\* Difference is significant at P value ≤0.05.
\*\* Difference is significant at P value ≤0.01.
P value calculated by Ordinal Logistic Regression (significant at P value ≤0.05).

In addition, the increased measures of obesity markers BMI and WC have been associated significantly with high hs-CRP tertile (P<0.01). BMI of low tertile (27.69±0.95 kg/m^2^) and moderate tertile (34.86±1.28 kg/m^2^) showed lower significance compared to high tertile (37.86±0.81 kg/m^2^); P<0.001 and P=0.011, respectively. Moreover, WC of low tertile had lower significance than high tertile (112.62±1.67 vs. 119.95±1.49 cm, P<0.009). Nevertheless, no difference was discovered for SBP, while DBP for high tertile had been higher than that with low significance (88.73±1.58 vs. 80.42±2.38 mmHg, P=0.026).

On the other hand, regarding metabolic markers, FBG, TC, and TG showed significant changes with differences in hs-CRP tertile. The lower tertile of hs-CRP had lower significance than higher category for FBG (146.90±12.92 vs. 207.33±9.64 mg/dL, P<0.001). As for TC, the lower tertile of hs-CRP (194.10±4.95 mg/dL) had lower significance than moderate (208.03±7.28 mg/dL) and high tertile (209.69±5.37 mg/dL); P=0.012 and P=0.008, respectively. Meanwhile, for TG, the lower tertile of hs-CRP (166.76±18.08 mg/dL) had lower significance than moderate (234.10±24.00 mg/dL) and high tertile (235.69±13.76 mg/dL); P=0.006 and P<0.001, respectively.

On the relation of adipokines, IL-6 and adiponectin showed significant changes with differences in hs-CRP tertile. For IL-6, the lower tertile of hs-CRP (1.60±0.10 pg/mL) had lower significance than moderate (2.05±0.12 pg/mL) and high tertile (2.39±0.11 pg/mL); P=0.005 and P<0.001, respectively. Inversely for adiponectin, the higher tertile of hs-CRP (8.55±0.53 mg/L) had lower significance than moderate (10.93±0.62 mg/L) and low tertile (13.20±0.98 mg/L); P=0.025 and P<0.001, respectively.

Therefore, in order to identify the risk factors associated with increasing a tertile of serum hs-CRP; Ordinal Logistic Regression was used after adjustments were made for age, gender, smoking habits, and physical activity *(Test of Parallel Lines:* P=0.968). However, the factors of WC, SBP, DBP, TC, and TG did not reach the significance level as risk factors. Nonetheless, the risk factors that had been associated with increased hs-CRP tertile were increased BMI [OR: 1.17, P=0.018], increased FBG [OR: 1.01, P=0.007], increased IL-6 [OR: 2.22, P=0.025], and reduced Adiponectin [OR: 0.81, P=0.002].

## DISCUSSION

This study identified the clinical risk factors associated with inflammation evidenced by high level of hs-CRP in hypertensive diabetic patients. CRP is a sensitive marker of systemic inflammation that is synthesized by the liver [37]. The elevated levels of CRP has been linked directly with the development of major adverse effects of cardiovascular diseases in T2DM patients [30], endothelial dysfunction and atherosclerosis [38], metabolic syndrome and T2DM [39], as well as HT in normotensive adults [40].

As reported by Den Hertog et al. [41], the increased levels of CRP had been associated with increased mortality and morbidity rates in patients with ischemic heart disease and stroke, as it resulted in poor outcome and death. In the present study, the higher tertile of hs-CRP had been associated with higher anthropometric measurements of BMI and WC. It was also associated with poor biomedical data represented by high blood pressure, dysglycemia, dyslipidemia, and disturbances in adipokines.

The significant increment of biomedical data with higher tertile of hs-CRP is consistent with that found in previous studies. For instance, Santos *et al.* [42] found that CRP levels were significantly higher in high BP, hypertriglyceridemia, and hyperglycemia in a representative sample of urban adults to evaluate the association between CRP and metabolic syndrome. Likewise, Kawamoto *et al.* [43] found that CRP was higher significantly among the FBG≥100 (mg/dL) group than the FBG<100 (mg/dL) group through a crosssectional study in Japan. In addition, Ajani *et al.* [44] found that the prevalence of high CRP increased for high lipid profile through an analysis of National Health and Nutrition Examination Survey 1999–2000.

Moreover, in order to identify the relationship of CRP with renal function loss in non-diabetic population, Stuveling *et al.* [45] found that the increased CRP had been associated with increased values of SBP, DBP, FBG, and TC. Similarly, Lee at al. [30] studied the role of inflammation with cardiovascular events, and discovered that CRP elevated significantly with the increase of DBP, FBG, TC, and TG, while the change of SBP did not reach the significance value with the change in CRP.

On the other hand, women in the present study had been associated with higher percentage of higher hs-CRP tertile, compared to men who achieved higher percentage of lower hs-CRP tertile. Consistently, in a metaanalysis conducted by Choi *et al.* [46], women had a stronger correlation of CRP than men in all ethnicity groups involving American, European, and Asian populations. Similarly, Thompson *et al.* [47] found that CRP was higher in women than men in China with high WC population, whereas Connelly *et al*. [48] agreed that the prevalence of high CRP (≥3.8 mg/L) among females was higher than the prevalence among male significantly in surveyed 512 community members aged 18 years and older in a study of Sandy Lake.

### Prediction of risk factors

In the Ordinal Logistic Regression analysis, the obtained risk factors that had been associated with increased hs-CRP were the increased BMI, FBG, and IL-6, as well as decreased adiponectin. Indeed, the relationship of the total obesity evidenced by high BMI with inflammation has been well-described [49]. Besides, obesity has the potential to generate oxidative stress and inflammation in a healthy population [50,51], and both of them could mediate the development of metabolic syndrome due to obesity [26,49].

Obesity is the main contributor to initiate the systemic inflammation in healthy adults though alteration on adipokine and/or inflammatory profiles are present [23]. Hansen et al. [52] compared the basal plasma adipokines and inflammatory markers between obese and non-obese T2DM by normoglycemic non-obese subjects and concluded that the observed alterations on adipokines and inflammatory markers in obese T2DM, as opposed to non-obese T2DM, had been attributed to the greater adipose tissue mass, and not necessarily due to the presence of diabetes.

Moreover, the overexpressed pro-inflammatory cytokines in obesity had been considered as the direct link between obesity and inflammation [51]. Adipose tissue responds to stimulation of extra nutrients via hyperplasia and hypertrophy of adipocytes [23]. The nature of adipose tissue is heterogeneous; including endothelium, immune cells, and adipocytes [53]. With progressive adipocyte enlargement and obesity, the blood supply to adipocytes is reduced and this leads to consequent hypoxia [54].

Hypoxia, on the other hand, has been proposed as an inciting etiology of necrosis and macrophage infiltration into adipose tissue, which leads to overproduction of pro-inflammatory mediators. This result in localized inflammation in adipose tissue that propagates an overall systemic inflammation associated with the development of obesity-related comorbidities [55]. Amongst the inflammatory mediators, three are produced by macrophages TNF-α, IL-6, and adiponectin [56]. After that, the produced IL-6 in the adipose tissue of healthy humans is released into the circulation; inducing systemic inflammation, which is evaluated by the sensitive marker synthesized primarily by the liver CRP [57]. Moreover, the accumulation of free fatty acids in obesity activates pro-inflammatory serine kinase cascades, such as I_k_B kinase and c-JunN-terminal kinase, which in turn, promotes adipose tissue to release IL-6 that triggers hepatocytes to synthesize and secrete CRP [58].

Other than that, the antioxidant defense factors become lower due to obesity and accumulation of fat [59], and due to obesity, the level of adiponectin, which is totally anti-inflammatory, is exhausted by the state of inflammation [60,61]. In several reports, adiponectin has been considered as an important regulator of insulin sensitivity and glucose homeostasis by confirming an inverse relationship with insulin resistance [62]. This inverse association may possibly be mediated not only by insulin, but also by inflammation induced by adipokines like IL-6 [63].

In this study, the results revealed that the increased BMI was a risk factor and a predictor of elevated hs-CRP. As shown in **Figure 1** (Chart A), there is a direct correlation between BMI and hs-CRP (*r*=0.345, P<0.001). As reported in previous studies, Lee et al. [30] found that the group with high tertile of hs-CRP was higher in BMI than other groups in a study intended to predict cardiovascular adverse effects in T2DM patients. Similarly, Pradhan et al. [64] evaluated the level of inflammatory markers among diabetic and healthy US women through a prospective case-control study to display the role of inflammatory markers CRP and IL-6. The level of CRP had a direct escalation by increasing the level of BMI for both case and control women. The mean of BMI for case group was (31.8 kg/m^2^) and for control group was (25.6 kg/m^2^); the median of CRP for case group was (0.69 mg/dL), which had been significantly higher than control group (0.26 mg/dL). Moreover, Escobar-Morreale et al. [65] concluded that obesity is the main determinant to elevate inflammatory markers CRP and IL-6 compared with lean bodies among pre-menopause women. In addition, Visser et al. [66] reported that the CRP level was higher among obese than non-obese in a study that included males and non-pregnant females aged 17 years and older.

**Figure 1.**
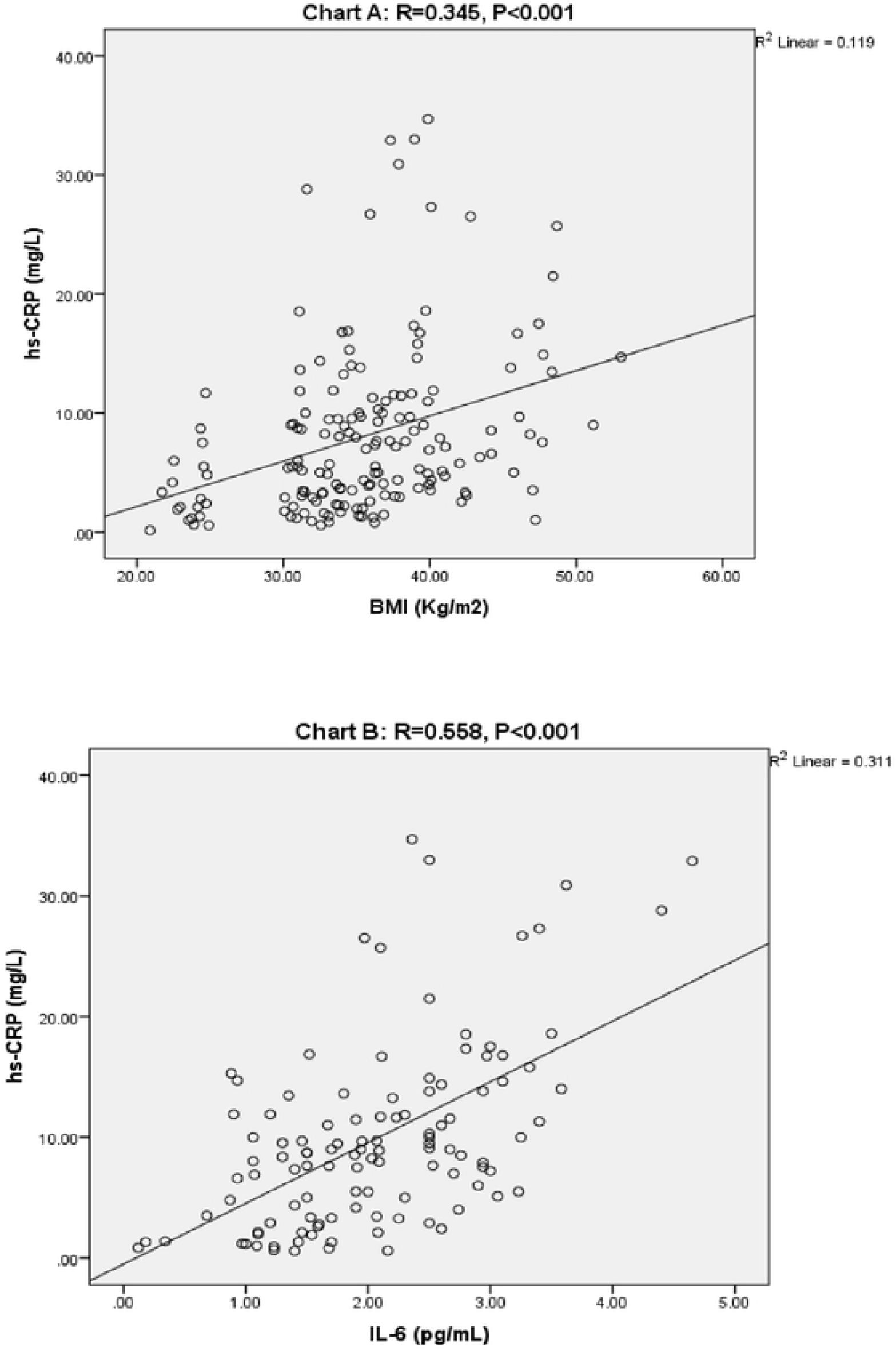
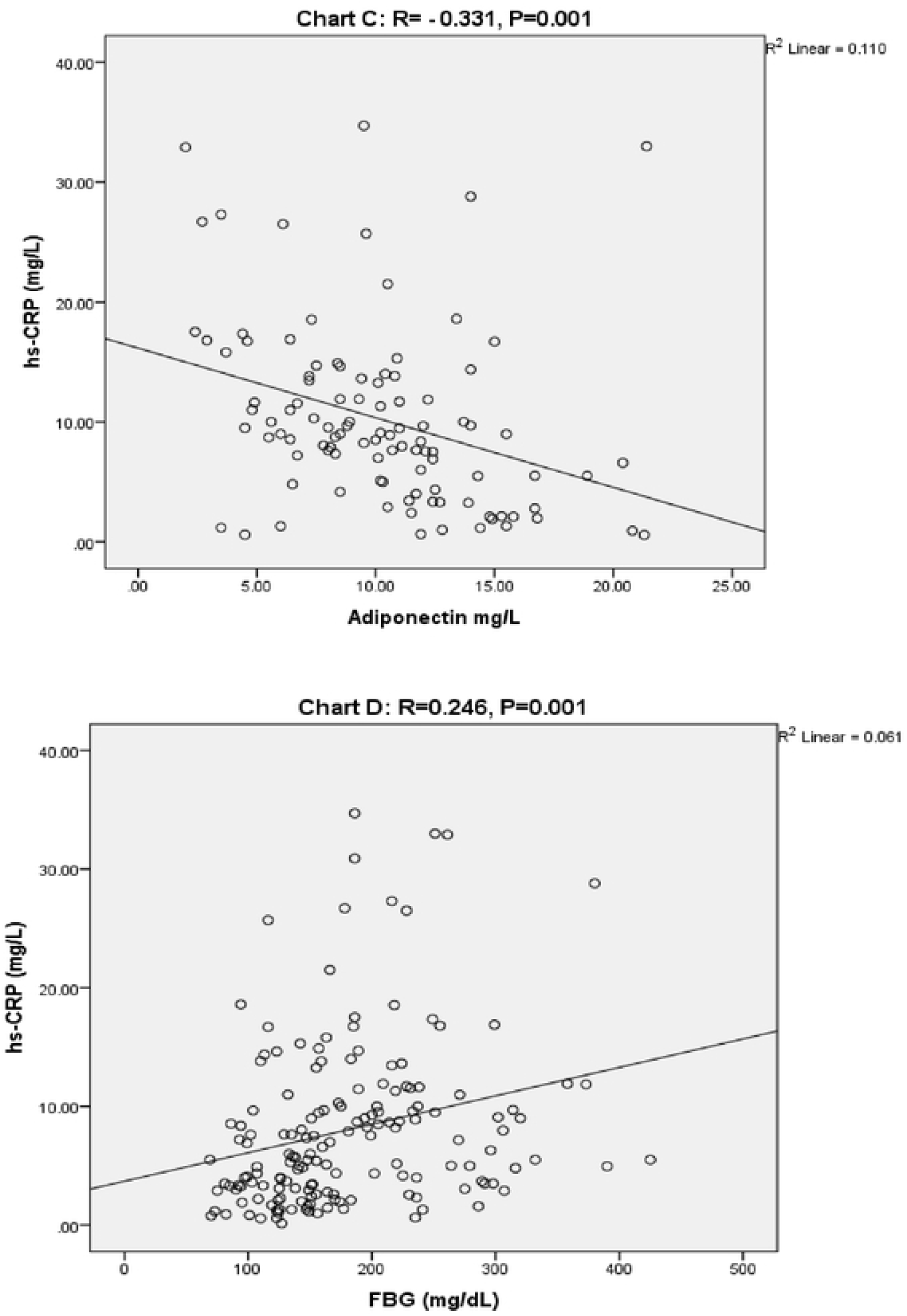
Bivariate correlations, Chart A: BMI with hs-CRP, Chart B: IL-6 with hs-CRP, Chart C: adiponectin with hs-CRP, Chart D: FBG with hs-CRP]

Furthermore, the results showed that the increased level of IL-6 was well-associated with elevated hs-CRP. As shown in **Figure 1** (Chart B), IL-6 directly correlated with hs-CRP (*r*=0.558, P<0.001). This result is strongly agreed by Indulekha *et al*. [67] who found that CRP increased in individuals who had elevated level of IL-6 among Asian Indians after distributing the population into four groups (diseased obese, healthy obese, diseased-non obese, and healthy non-obese). Similarly, Wannamethee *et al*. [68] found a strong and significant association between IL-6 and CRP (P<0.001) during the evaluation of risk factors associated with metabolic syndrome and CVDs among overaged British in several towns. In the same vein, Tsilchorozidou *et al.* [69] found strong correlation between IL-6 and CRP (*r*= 0.59) in women with polycystic ovary syndrome. In addition, Ridker *et al*. [70] studied the association between IL-6 and CRP in 202 myocardial infarction patients and 202 apparently healthy participants matched by age and sex; where significant correlation was indicated (*r*=0.43, P<0.001). In fact, the results of the current study and past literatures have approved the strong association and effect of IL-6 on the elevation of hs-CRP.

Moreover, the results represented that the decreased levels of adiponectin were well-associated with elevated hs-CRP. As shown in **Figure 1** (Chart C), adiponectin inversely correlated with hs-CRP (*r*= −0.331, P=0.001). This result is consistent with that obtained by Sedighi and Abediankenari [71], in a study on chronic kidney disease patients, where they found inverse correlation between adiponectin and CRP (*r*= – 0. 548, P< 0.001). Likewise, Hung *et al.* [72] found that the levels of CRP and IL-6 could predict negative values of adiponectin after adjustments were to age, sex, and waist to hip ratio; where the estimated regression coefficient was [−4.56], P<0.001, and [−3.46], P=0.001, respectively, while the estimation of regression was still significant after adding smoking habits.

Finally, this study proved the effect of diabetes, evidenced by high level of FBG on inflammation featured by hs-CRP. This effect has further asserted the positive correlation between FBG and hs-CRP (*r*=0.246, P=0.001), as shown in **Figure 1** (Chart D). Similar results were also observed in previous studies. Kawamoto *et al.* [43] found CRP higher significantly (P=0.033) among the FBG≥100 (mg/dL) group than the FBG<100 (mg/dL) group through a cross-sectional study carried out in Japan. Likewise, Santos *et al.* [42] found that the CRP levels were significantly higher among urban adults with high FBG compared to low level (1.96 vs. 1.46, P=0.032) in order to evaluate the association between CRP and metabolic syndrome.

## CONCLUSION

The presence of inflammation featured by hs-CRP could predict metabolic dysregulation in hypertensive patients with T2DM. The group of patients with higher tertile of hs-CRP displayed major metabolic abnormalities, including dysglycaemia, dyslipidemia, higher BP, and metabolic obesity indexed by adipokines activity. Besides, the results of the investigation retrieved from this study proved the role of obesity in promoting systemic inflammation. BMI was a risk factor of elevated hs-CRP, while the conjugated risk factors of increased IL-6 and reduced adiponectin were strongly linked to obesity [23,56]. In fact, similarity in these results was observed in Al-Hamodi et al. [27] study that proved the role of obesity on adiponectin and inflammation. In addition to obesity, this study proved the role of diabetes evidenced by hyperglycemia in initiating inflammation that could lead to major adverse effects. Therefore, controlling diabetes and managing weight through positive lifestyle patterns can reduce the incidence of chronic inflammation and further complications.

## ACKNOWLEDGEMENT

The authors would like to extend their sincere gratitude to the participants for their willingness to participate in this study, and to the Palestinian Ministry of Health for granting the permission to conduct the fieldwork. In addition, the author would also like to thank the Faculty of Medicine and Health Sciences at Universiti Putra Malaysia for the use of its library.

## CONFLICT OF INTEREST

The author declare that there is no significant competing financial, professional or personal interests that may influence the performance or the presentation of the work described in this manuscript.

